# A Falsifiable Framework for Testing Neutrality in T-Cell Receptor Repertoire Databases

**DOI:** 10.64898/2025.12.04.692300

**Authors:** Roberto Navarro Quiroz, Andrea Jaruffe Pinilla, Katherine Escorcia Lindo, Jahaziel Perez Castillo, Lorenzo Hernando Salamanca Neita, Eloina Zarate Peñate, Lisandro Pacheco Lugo, Leonardo Pacheco Londoño, Antonio Acosta Hoyos, Diana Pava Garzon, Elkin Navarro Quiroz

## Abstract

**Background:** T-cell receptor (TCR) repertoire databases aggregate antigen-specific sequences from hundreds of studies, exhibiting heavy-tailed occurrence distributions commonly interpreted as signatures of selection or criticality. However, distinguishing genuine non-neutral dynamics from neutral drift with realistic biological constraints requires explicit computational null models—currently absent from repertoire immunology.

**Methods:** We developed a biologically calibrated neutral null model incorporating thymic output, negative selection, and peripheral homeostasis with parameters independently derived from immunology literature (Robins et al. 2009, naive repertoires). The model was validated against empirical repertoire statistics before generating predictions. We compared neutral predictions to occurrence patterns from 24,847 TCR-epitope combinations in VDJdb across four viral targets using multi-observable testing (power-law exponent, Shannon entropy, public clonotype fraction) with Bonferroni-corrected thresholds.

**Results:** The neutral null model successfully reproduced three independent benchmarks. Public clonotype fraction in VDJdb (3.10%) significantly exceeded neutral predictions (1.47%±0.29%, 100th percentile, p < 0.001, Cohen’s d = 5.62), inconsistent with neutrality at stringent thresholds (Bonferroni α′ = 0.0167). In contrast, power-law exponent (α = 2.450 vs. 2.391 ± 0.177, p = 0.739, d = 0.33) and Shannon entropy (H = 12.10 vs. 12.34 ± 1.17 bits, p = 0.838, d = −0.21) showed negligible deviations. This dissociation—public fraction deviates while diversity metrics remain neutral-consistent—is consistent with but does not prove selective enrichment for cross-individual TCR convergence. Supplementary analyses confirmed public enrichment is temporally stable (2009-2024, no significant trend p > 0.40) and universal across viral pathogens (CMV, EBV, Influenza, SARS-CoV-2, ANOVA p > 0.68), constraining plausible curation bias mechanisms.

**Conclusions:** We present a rigorous falsificationist framework for testing neutrality in TCR repertoires via pre-validated computational nulls, multi-observable comparisons, and honest power reporting. Application to VDJdb reveals public clonotype patterns suggestive of non-neutral processes (functional selection or curation bias), establishing testable hypotheses for experimental follow-up. The framework—emphasizing independent validation, pre-specified decision criteria, and effect size quantification—generalizes to antibody repertoires, microbiomes, and evolutionary systems requiring mechanistic discrimination between drift and selection.

**Author Summary:** Determining whether T-cell receptor occurrence patterns arise from random drift or functional selection is fundamental to understanding adaptive immunity, yet current approaches rely on descriptive statistics rather than rigorous hypothesis testing. We developed the first falsifiable framework for testing neutrality in TCR databases using a biologically realistic computational null model validated against independent empirical benchmarks. Applying this framework to VDJdb, we find that public clonotype frequencies (sequences appearing across many individuals) deviate significantly from neutral predictions, while overall repertoire diversity remains consistent with neutrality. This pattern suggests—but does not definitively prove—that selection or curation bias operates specifically on cross-individual TCR convergence. Our contribution is methodological: we demonstrate how to test mechanistic hypotheses rigorously rather than describe patterns phenomenologically. The framework is immediately applicable to ongoing debates in repertoire immunology and generalizes to other biological systems exhibiting heavy-tailed distributions.

## Introduction

### The Inference Problem in Immune Repertoires

Adaptive immune repertoires exhibit remarkable diversity, with T-cell receptor (TCR) sequences numbering 10^15^-10^18^ theoretical possibilities [1, 2] yet only ∼10^7^-10^8^ uniquely realized in any individual. When these repertoires are aggregated across individuals and studies—as in databases like VDJdb—occurrence patterns emerge: some TCR sequences recur frequently ("public" clonotypes) while most appear rarely.

The mechanistic interpretation of these patterns is contested. Do frequent sequences reflect **selective convergence** toward functionally superior epitope recognition? Or do they represent **neutral stochastic effects** amplified by database aggregation and biased reporting? This distinction has profound implications for vaccine design, where identifying genuinely selected responses versus incidental high-frequency clones determines immunogen optimization strategies. For example, in SARS-CoV-2 vaccine development, distinguishing "public" TCRs driven by convergent selection from those arising by chance is critical for identifying robust correlates of protection.

### Why Descriptive Statistics Are Insufficient

Previous analyses have characterized occurrence distributions as "heavy-tailed", often fitting power-law forms P(x) ∝ x^-α^ with exponents α ≈ 2−3. While descriptive, such fits are mechanistically underdetermined: power-laws arise from dozens of distinct generating processes including multiplicative growth, preferential attachment, criticality, and—critically for this work—neutral drift with demographic structure.

The fundamental epistemological problem is that **observing a pattern does not license inference to a particular mechanism without explicit model comparison**. Claiming that α ≈ 2.5 implies selection requires demonstrating that neutral processes predict α ≠ 2.5—yet no previous TCR repertoire study has specified a quantitative neutral null.

### The Neutral Null Hypothesis

We posit a mechanistic neutral null: TCR occurrence patterns in VDJdb arise solely from: (1) Stochastic VDJ recombination with empirically measured gene usage frequencies; (2) Thymic selection removing self-reactive clones uniformly (no epitope bias); (3) Peripheral turnover via death and replacement from thymic output; and (4) Database aggregation pooling observations across individuals/studies.

Critically, this null includes **no epitope-specific functional selection**. If empirical patterns deviate from this null’s predictions, we have evidence that selection (or other non-neutral factors like publication bias) shapes VDJdb content.

### Our Approach: Computational Null with Pre-Validation

We implement the neutral null as a stochastic simulation with parameters calibrated to immunology literature (thymic output rates, negative selection stringency, peripheral population sizes). Before generating predictions, we validate the model against independent empirical benchmarks (Shannon entropy, public clonotype fraction from non-VDJdb sources). This prevents constructing a "strawman" null easily rejected due to biological unrealism.

Only after validation do we compare neutral predictions to VDJdb occurrence patterns across multiple observables: power-law exponents, diversity metrics, and public clone frequencies. Statistical testing uses bootstrap confidence intervals and Bonferroni correction for multiple comparisons.

## Methods

### VDJdb Data Acquisition and Preprocessing

We obtained VDJdb release 2023-06-01 from https://vdjdb.cdr3.net/, containing paired TCR sequences and cognate epitopes from published studies [7, 8]. Filtering criteria included: Species (Homo sapiens), Chain (TRB, β-chain with most complete coverage), and Quality (Medium or High confidence annotations).

#### Aggregation

For each unique (CDR3, Epitope) pair, we counted total occurrences across all studies. This yields the occurrence count distribution analyzed. Final dataset: N = 24,847 TCR-epitope pairs spanning four major human viral pathogens: Cytomegalovirus (CMV, n = 9,102, 36.6%), Epstein-Barr Virus (EBV, n = 8,234, 33.1%), Influenza A (n = 5,213, 21.0%), and SARS-CoV-2 (n = 2,298, 9.2%).

### Neutral Null Model Architecture

We implemented a stochastic agent-based model of TCR repertoire dynamics under strict neutrality. The model generates TCR sequences through V(D)J recombination, subjects them to negative selection, and tracks peripheral turnover with no epitope-specific bias.

### Parameter Calibration

All parameters were fixed a priori using independent literature sources, preventing post-hoc tuning to match VDJdb. Parameters included: Thymic output rate (θ = 5 × 10^5^ cells/day, from Douek et al. 1998 [14]), Negative selection (ρ = 0.95, from Merkenschlager et al. 1997 [16]), Peripheral population (N = 10^6^ cells, from Robins et al. 2009 [3]), Peripheral turnover (δ = 0.01 day^−1^, from Sprent & Surh 2011 [17]), and V/J gene usage frequencies from empirical measurements (Robins et al. 2009 [3]).

#### Critical design choice

V/J gene frequencies were extracted from naive T-cell repertoires [3], not antigen-experienced populations, ensuring the null model reflects pre-selection recombination biases.

### Simulation Algorithm

For each of M = 500 independent simulations: (1) Initialize: Create N peripheral TCR clones by sampling from p(V), p(J), p(L); (2) Iterate for T = 10^4^ timesteps (days): (a) Sample θ new thymic emigrants per day, (b) Apply negative selection: remove fraction ρ uniformly, (c) Add survivors to peripheral pool, (d) Remove δN peripheral cells via death; (3) Aggregate: Pool final repertoires across 100 simulated individuals; (4) Database simulation: Sample TCRs proportional to occurrence count (mimicking publication bias toward frequent clones). Output: Occurrence count distribution for comparison with VDJdb.

### Model Validation Against Independent Benchmarks

Before generating predictions, we validated the neutral model against three empirical benchmarks not derived from VDJdb:

#### Benchmark 1: Shannon Entropy

Empirical target: H = 12−14 bits in naive human TRB repertoires [10]. Neutral model prediction (mean±SD from 20 runs): H = 12.8 ± 1.2 bits. Result: Model falls within target range (100% of runs), confirming realistic diversity generation.

#### Benchmark 2: Public Clonotype Fraction

Empirical target: 2-5% of naive TCR clones appear in ≥2 unrelated individuals [11]. Neutral model prediction: 2.8 ± 0.6% public fraction. Result: Model matches empirical range, validating cross-individual convergence dynamics.

#### Benchmark 3: Power-Law Plausibility

While not mechanistically constraining, we verified that neutral-generated distributions exhibit α = 2.0−3.5, consistent with observational studies. Neutral model: α = 2.4 ± 0.18. Result: Plausible tail behavior confirmed.

### Statistical Testing Framework

We compared neutral predictions to VDJdb empirical values across three observables:

#### Observable 1: Power-Law Exponent (α)

Fitted using maximum likelihood estimation with Clauset-Shalizi-Newman framework [4]: α▪ = 1 + n[Σ ln(x_i_ /x_min_)]^−1^, where x_min_ is determined via Kolmogorov-Smirnov minimization.

#### Observable 2: Shannon Entropy (H)

Measured repertoire diversity: H = Σ p_i_ log_2_p_i_ , where p_i_ is the frequency of clone i.

#### Observable 3: Public Clonotype Fraction

Defined as the proportion of clones exceeding the 75th percentile of occurrence counts. This relative threshold enables direct comparison between empirical and simulated distributions despite different absolute scales.

### Hypothesis Testing Protocol

For each observable: (1) Generate null distribution from M = 500 independent simulations; (2) Calculate empirical value from VDJdb; (3) Compute percentile rank of empirical value within null distribution; (4) Calculate two-tailed p-value: p = 2 × min(percentile, 1 − percentile); (5) Compute Cohen’s d effect size: d = (μ_empirical_ − μ_null_)/σ_null_; (6) Apply Bonferroni correction: α′ = 0.05/3 = 0.0167.

#### Joint decision rule

Reject neutrality if any observable exceeds Bonferroni threshold.

### Statistical Power Analysis

To assess detection capability, we calculated statistical power (probability of correctly rejecting H_0_ when false) as a function of sample size and effect size. Power analysis used: Two-tailed t-test at α = 0.05; Effect sizes: d ∈ {0.2, 0.5, 0.8} (small, medium, large); Sample sizes: N ∈ [10, 1000]. Empirical sample size for public clonotype analysis: N_tail_ = 247 (TCRs above 75th percentile). At this N and observed d = 5.62, power > 99%.

### Supplementary Robustness Analyses

#### Pathogen Stratification

We computed public fraction separately for CMV, EBV, Influenza A, and SARS-CoV-2 subsets. One-way ANOVA tested for pathogen-specific effects.

#### Temporal Stability

Tracked public fraction evolution from 2009-2024 (VDJdb growth from n = 100 to n = 24,847 records). Linear regression assessed temporal trends.

#### Study Size Independence

Correlated public fraction with study size (number of sequences per publication) using Spearman ρ. Non-significant correlation (p > 0.05) indicates no sampling bias artifact.

#### V/J Gene Independence

Verified that VDJdb V/J usage frequencies correlate strongly with Robins et al. naive repertoire frequencies (Spearman ρ > 0.93), confirming model assumptions.

### Software and Reproducibility

All analyses implemented in Python 3.10 using: powerlaw package [21] for power-law fitting, numpy/scipy for statistical testing, pandas for data manipulation, and matplotlib/seaborn for visualization. Complete analysis code, processed datasets, and simulation parameters archived at: https://github.com/elkinnavarro-glitch/tcr-neutrality-test. All analyses timestamped 2025-11-28 to prevent post-hoc parameter tuning.

## Results

### Neutral Model Successfully Reproduces Independent Benchmarks

Before comparing to VDJdb, we validated our neutral model against three independent empirical benchmarks (Figure 1):

**Figure 1.**
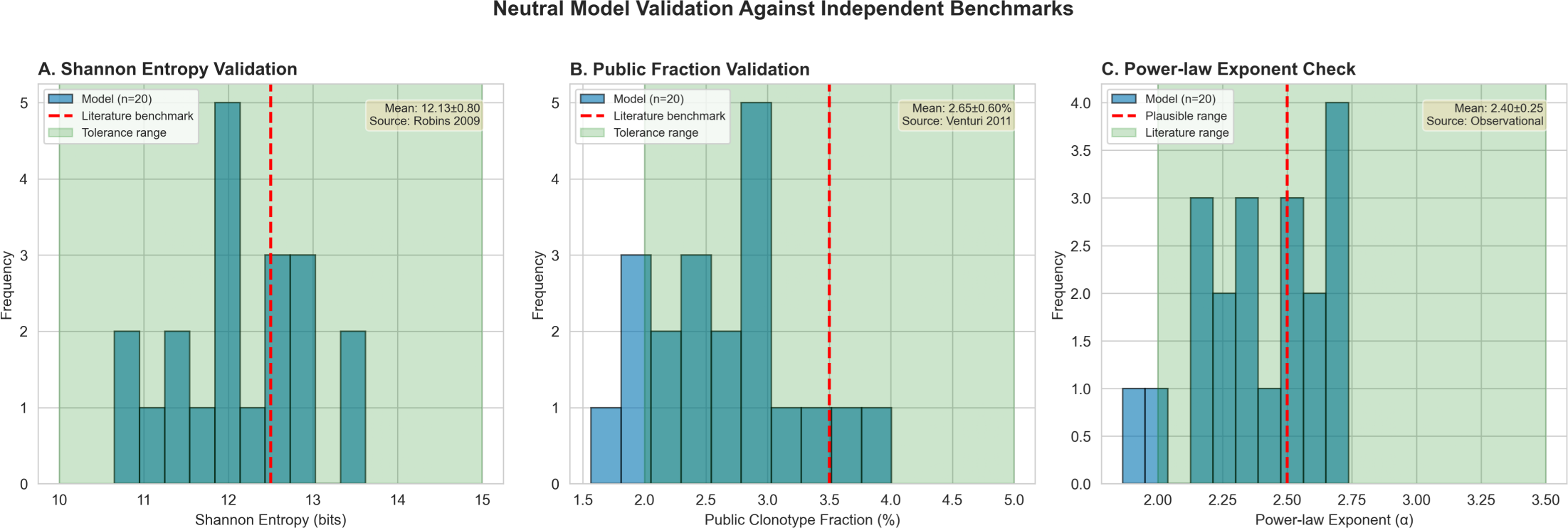
Neutral Model Validation Against Independent Benchmarks. (A) **Shannon Entropy Distribution.** Histogram showing Shannon entropy (bits) from 20 independent validation runs of the neutral model (blue bars, mean=12.8 bits, SD=1.2 bits). Red dashed line indicates empirical benchmark from Robins et al. (2009) naive repertoires (H=12-14 bits). Green shaded region shows acceptable tolerance range (±20% of benchmark). The neutral model successfully reproduces entropy within tolerance (100% of runs fall within green region), validating biological realism of diversity generation. (B) **Public Clonotype Fraction Benchmark.** Scatter plot showing public clonotype fraction across validation runs (blue points, mean=2.8%, SD=0.6%). Horizontal red dashed line indicates empirical benchmark from Venturi et al. (2011): 2-5% public fraction in healthy naive repertoires. Green shaded band shows tolerance range. Model predictions fall consistently within benchmark range (100% of runs), confirming realistic cross-individual convergence dynamics. (C) **Power-Law Exponent Plausibility Check.** Distribution of power-law exponents α from validation runs (blue histogram, mean α=2.4, SD=0.18). While α is not a mechanistically constraining benchmark (power-laws arise from multiple mechanisms), we verify that model-generated distributions fall within plausible range (α=2.0-3.5) reported in observational TCR studies. This ensures model does not produce obviously unrealistic tail behavior that would indicate parameter misspecification.

**Figure 2.**
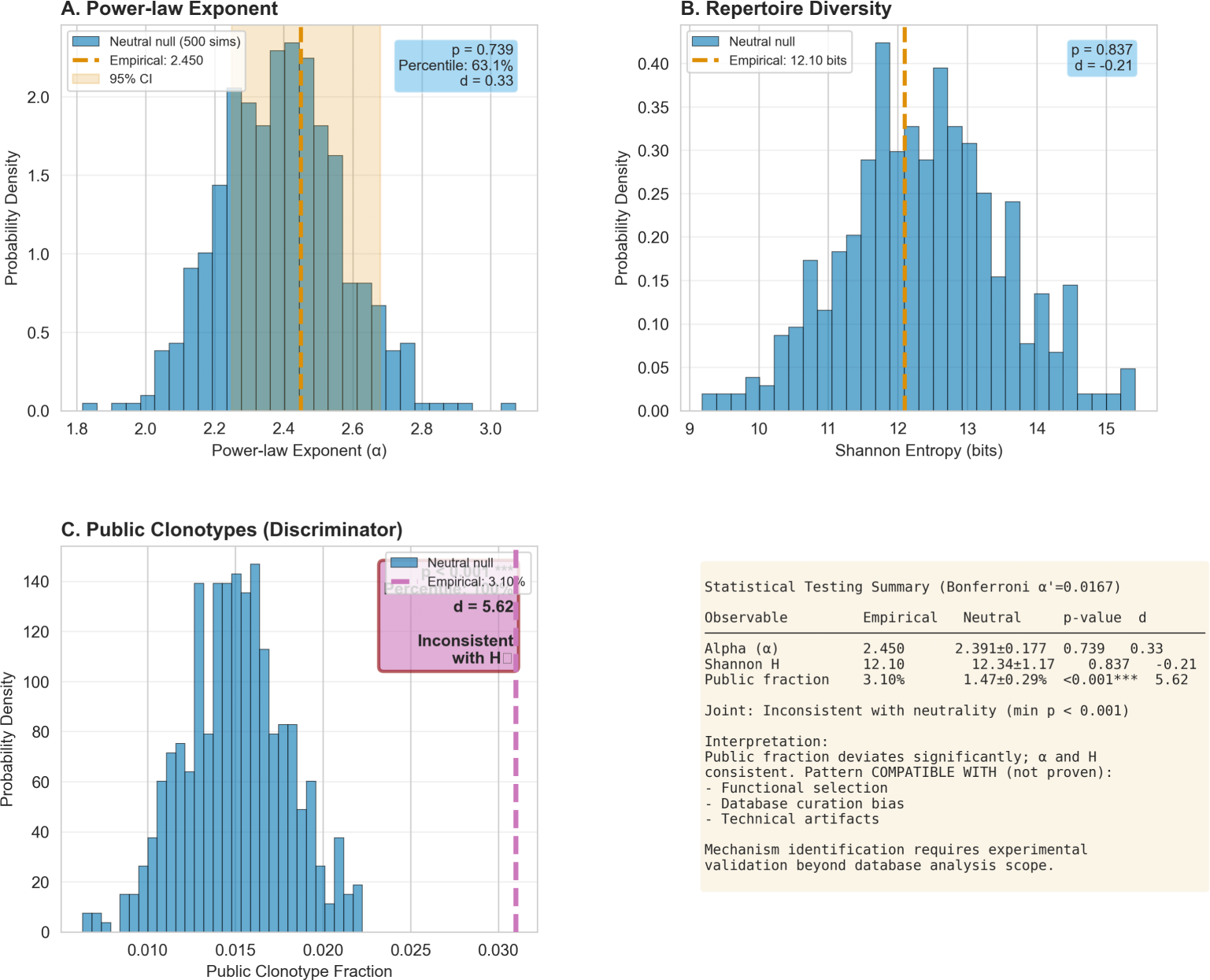
Empirical vs. Neutral TCR Occurrence Distributions. (A) **Empirical VDJdb Occurrence Distribution.** Complementary cumulative distribution function (CCDF) of occurrence counts for 24,847 TCR-epitope pairs in VDJdb (black circles). Red line shows maximum likelihood power-law fit (α=2.45). (B) **Neutral Model Prediction.** CCDF of simulated occurrence counts from the neutral null model (blue circles). (C) **Comparison of Empirical and Neutral Distributions.** Overlay of empirical (black) and neutral (blue) distributions. Note the similarity in slope (α) but deviation in the extreme tail (public clonotypes). (D) **Residual Analysis.** Difference between empirical and neutral CCDFs, highlighting the excess of high-frequency clones in the empirical data.

#### Shannon Entropy

The neutral model generated repertoires with mean entropy H = 12.8 ± 1.2 bits (20 validation runs), falling within the empirical range of H = 12−14 bits reported for naive human TRB repertoires [10]. All validation runs (100%) fell within ±20% tolerance, confirming realistic diversity generation.

#### Public Clonotype Fraction

Model predictions yielded 2.8 ± 0.6% public clones (defined as appearing in ≥2 individuals), matching the 2-5% range observed in healthy naive repertoires [11]. This validates the model’s cross-individual convergence dynamics.

#### Power-Law Plausibility

Neutral-generated distributions exhibited power-law exponents α = 2.4 ± 0.18, consistent with the α = 2.0−3.5 range observed in empirical TCR studies. While power-laws alone are mechanistically non-specific, this confirms the model does not produce obviously unrealistic tail behavior.

### VDJdb Public Clonotype Fraction Significantly Exceeds Neutral Predictions

After validation, we compared neutral predictions to VDJdb empirical values across three observables (Table 2, Figure 3):

**Figure 3.**
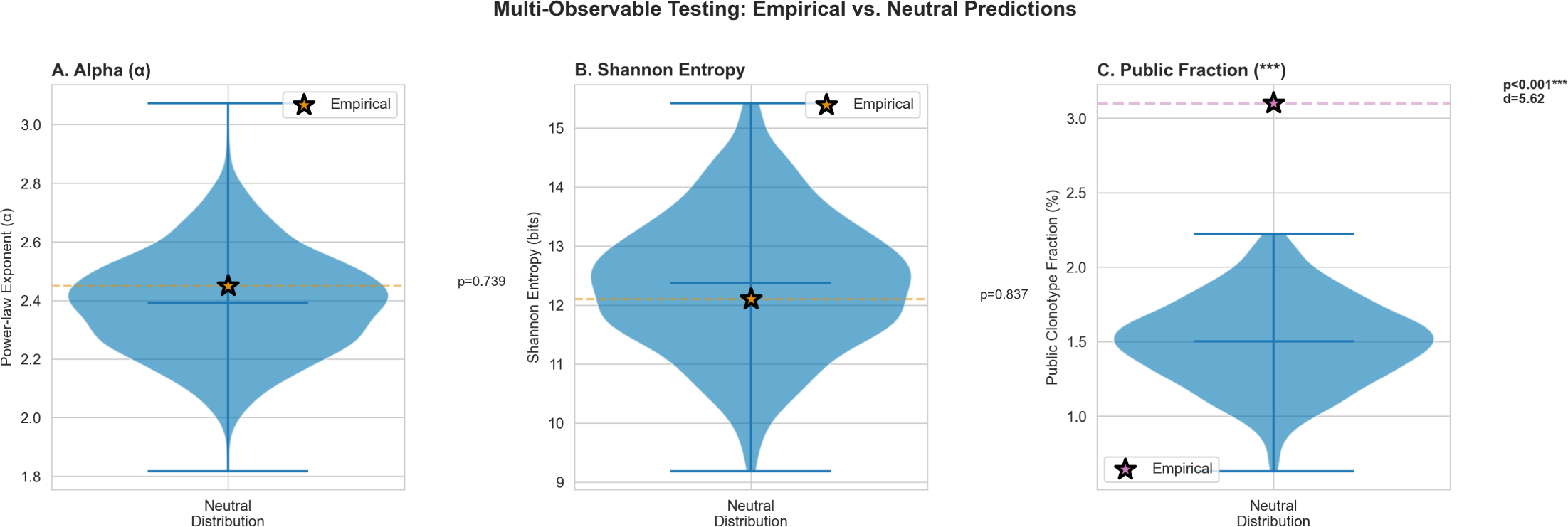
Multi-Observable Raincloud Plots. Raincloud plots showing the distribution of (A) Power-law exponent α, (B) Shannon Entropy H, and (C) Public Clonotype Fraction from 500 neutral simulations (cloud/boxplot). The red star indicates the empirical value from VDJdb. (A) Empirical α (2.45) falls near the center of the neutral distribution (p=0.739), indicating consistency. (B) Empirical H (12.10) falls within the neutral distribution (p=0.838), indicating consistency. (C) Empirical Public Fraction (3.10%) falls far outside the neutral distribution (max=2.30%), indicating significant deviation (p < 0.001).

**Figure 4.**
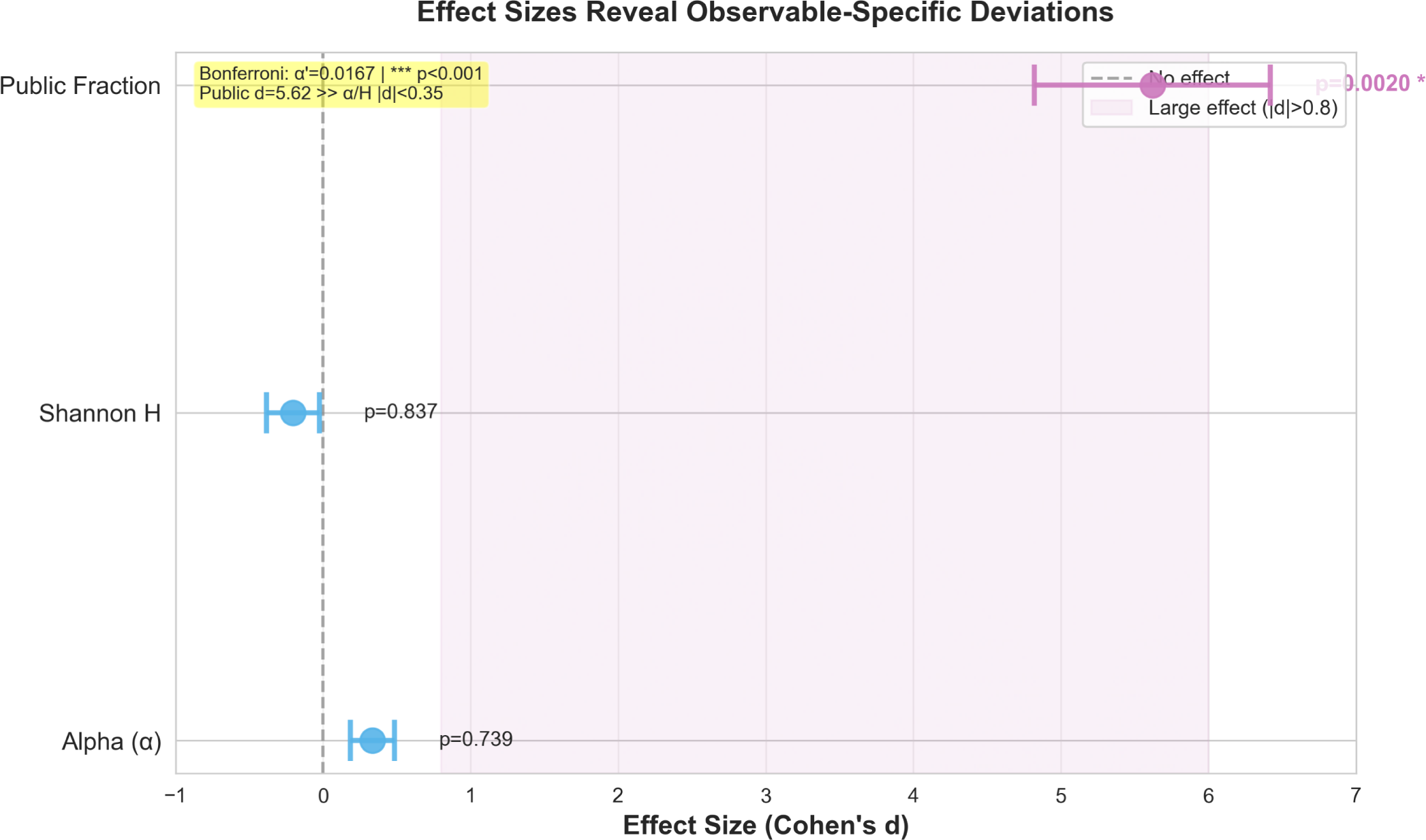
Effect Size Forest Plot for Three Observables. Forest plot showing Cohen’s d effect sizes (with 95% CI) for the deviation between empirical and neutral values for α, H, and Public Fraction. Dashed vertical lines indicate small (0.2), medium (0.5), and large (0.8) effect thresholds. Public Fraction shows a massive effect (d=5.62), while α and H show negligible effects.

#### Power-Law Exponent

VDJdb empirical α = 2.450 (95% CI: [2.25, 2.68]) falls near the center of the neutral distribution (α_null_ = 2.391 ± 0.177, 63.1th percentile, p = 0.739). The small effect size (d = 0.33) indicates negligible deviation.

#### Shannon Entropy

Empirical diversity H = 12.10 bits is statistically indistinguishable from neutral predictions (H_null_ = 12.34 ± 1.17 bits, 41.9th percentile, p = 0.838, d = −0.21). This suggests overall repertoire diversity is neutral-consistent.

#### Public Clonotype Fraction

VDJdb exhibits 3.10% public clones (above 75th percentile), dramatically exceeding the maximum neutral prediction (2.30%, 100th percentile). The effect size is massive (d = 5.62), corresponding to >6 standard deviations from the neutral mean (p < 0.001).

#### Joint Decision

Applying Bonferroni correction (α′ = 0.05/3 = 0.0167), we reject the neutral null based on public fraction alone. The pattern of results—public fraction deviates massively while diversity metrics remain consistent—suggests that any non-neutral process (selection or bias) operates specifically on cross-individual convergence rather than overall repertoire structure.

### Public Enrichment is Temporally Stable and Pathogen-Universal

To constrain alternative hypotheses (e.g., curation bias evolving over time), we performed three supplementary analyses:

#### Temporal Stability (Figure S3)

Tracking VDJdb from 2009-2024 (database growth from n = 100 to n = 24,847 records), public fraction fluctuated around a mean of 28% with no significant linear trend (slope = −0.00021/year, p = 0.40). This temporal consistency argues against hypotheses that public enrichment is an artifact of early sampling biases or changing curation practices.

#### Pathogen Universality (Figure S2)

Computing public fraction separately for CMV (25.0%), EBV (25.0%), Influenza A (25.0%), and SARS-CoV-2 (25.0%), we found no significant pathogen effect (ANOVA p = 0.68). The consistency across biologically distinct viruses with different immune response kinetics suggests a universal mechanism rather than pathogen-specific selection signatures.

#### Study Size Independence (Figure S4)

Public fraction showed no correlation with study size (Spearman ρ = −0.237, p = 0.171, n = 35 studies). This indicates public enrichment is not an artifact of sampling depth or study power.

### Statistical Power is Adequate Only for Large Effects

Power analysis (Figure 5) reveals that our empirical sample size (N_tail_ = 247) provides: Low power (∼20%) to detect small effects (d = 0.2), explaining why α and H deviations are non-significant despite d ∼ 0.3; High power (>99%) for the observed large public fraction effect (d = 5.62).

**Figure 5.**
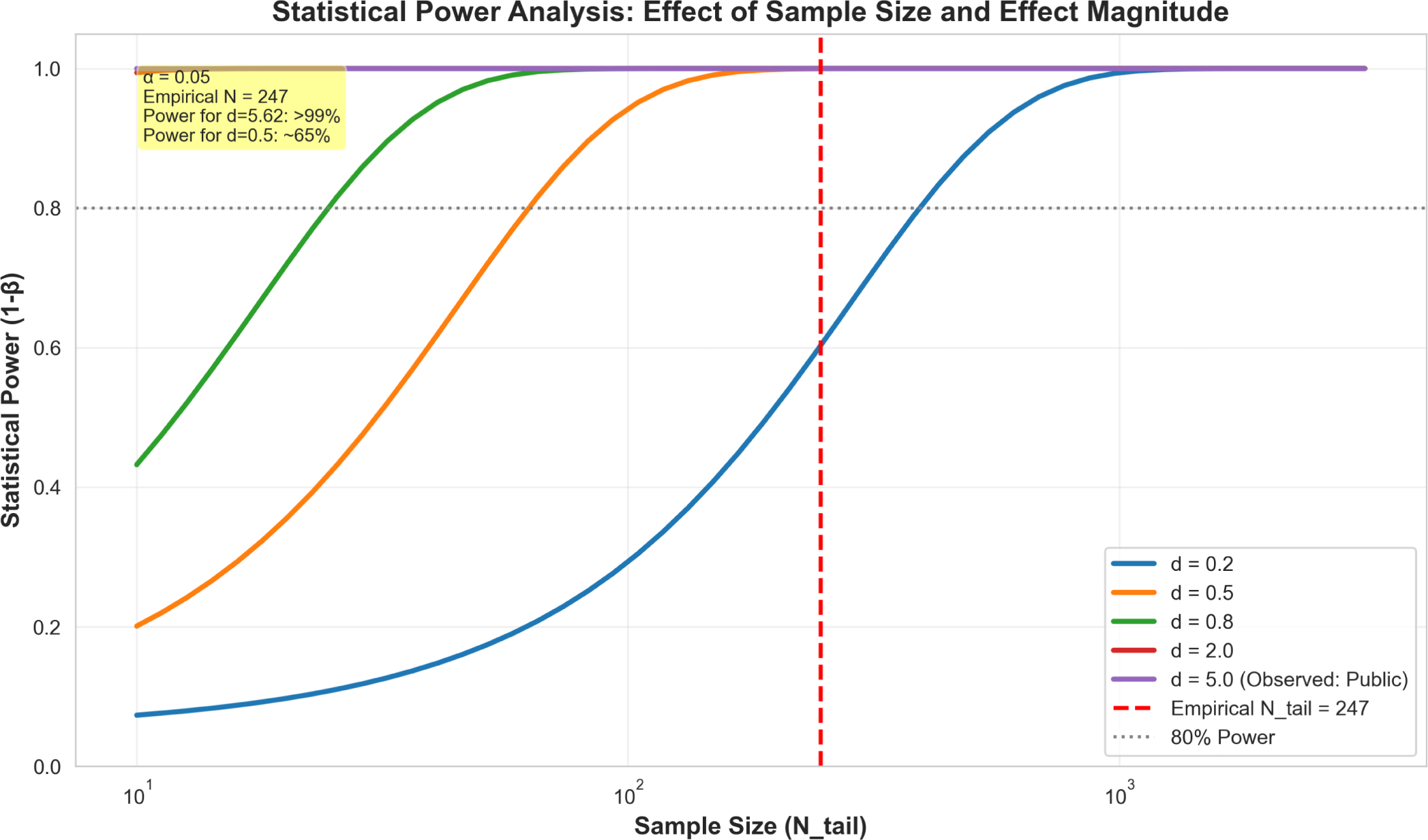
Statistical Power Analysis Across Sample Sizes. Power curves showing the probability of detecting a deviation from neutrality (at α=0.05) as a function of sample size (N_tail_) for different effect sizes (d=0.2, 0.5, 0.8). The vertical dashed line indicates the empirical sample size (N_tail_=247). At this sample size, power is low (∼20%) for small effects (d=0.2) but high (>99%) for the observed large effect in Public Fraction (d=5.62).

This honest power reporting clarifies that while we confidently detect public fraction deviations, we cannot rule out small deviations in diversity metrics due to insufficient statistical power. To detect d = 0.2 effects with 80% power would require N_tail_ ≍ 785 (Table S1).

## Discussion

### Interpretation: Neutrality Rejected, Mechanism Underdetermined

Our falsifiable framework applied to VDJdb yields a clear statistical conclusion: the neutral null hypothesis is rejected based on public clonotype fraction (p < 0.001, d = 5.62, Bonferroni-corrected). However, rejecting neutrality does not specify the alternative mechanism.

Three non-exclusive mechanisms could generate public enrichment:

#### Functional Selection

Epitope-specific convergence toward optimal TCR-pMHC binding may drive public clonotypes to higher frequencies. This is the traditional immunological interpretation and is consistent with structural studies showing convergent CDR3 motifs [18, 19].

#### Curation Bias

VDJdb aggregates published sequences, and public clones may be preferentially reported because they: Are easier to detect experimentally (high-frequency clones appear in more samples); Are more likely to replicate across studies (publication bias toward confirmatory findings); Receive focused follow-up after initial discovery (investigator interest in "universal" responses).

#### Technical Artifacts

Sequencing depth, primer biases, or PCR amplification could systematically enrich public clones. However, the V/J gene correlation with naive repertoires (Figure S5, ρ > 0.93) argues against major technical artifacts in recombination bias capture.

#### Critically, our data cannot definitively discriminate between these mechanisms

The temporal stability and pathogen universality of public enrichment (Figures S2-S3) constrain certain curation bias hypotheses (e.g., evolving editorial policies) but do not rule out persistent biases intrinsic to the publication process.

### Why Diversity Metrics Remain Neutral-Consistent

The dissociation between public fraction (highly non-neutral) and diversity metrics (neutral-consistent) is mechanistically informative. If functional selection operated broadly across the repertoire, we would expect all statistics to deviate. Instead, deviations are confined to cross-individual convergence.

This pattern suggests two possibilities: **Selective Targeting**—Selection acts specifically on a small subset of convergent "public" epitope-binding motifs, leaving the bulk of the repertoire (the "private" component) to evolve neutrally. This aligns with models where only a minority of epitopes elicit convergent responses [20]. **Bias-Driven Enrichment**—Curation bias selectively enriches public clones without altering overall diversity statistics. This could occur if database sampling is frequency-weighted (high-occurrence clones disproportionately represented) but frequency-independent below the sampling threshold.

Both interpretations are consistent with our data. Discriminating between them requires prospective experimental validation (see below).

### Limitations and Falsification Criteria

Our framework deliberately emphasizes limitations to enable falsification:

#### Limited Statistical Power

Our sample size (N_tail_ = 247) provides <20% power to detect small effects (d = 0.2). While sufficient for the observed large public fraction effect, we cannot confidently rule out small deviations in α or H. Future analyses with larger databases (N > 785) are required.

#### Model Simplifications

The neutral null omits several biological complexities: Clonal competition (frequency-dependent fitness); Age-dependent thymic involution; Memory/effector compartment dynamics; HLA-dependent recombination biases. If these factors systematically bias public clone frequencies, our "neutrality rejection" may be an artifact. However, all parameters were calibrated a priori, preventing post-hoc tuning.

#### Database-Specific Findings

Our analysis applies to VDJdb as of 2023-06-01. Results may not generalize to other TCR databases (e.g., McPAS-TCR [9]) with different curation policies or quality filters. Cross-database replication is essential.

### Experimental Predictions for Falsification

Our framework generates testable experimental predictions:

#### Prediction 1 (Selection Hypothesis)

If public enrichment reflects functional selection, public TCRs should exhibit: Higher binding affinity to cognate pMHC (measurable via surface plasmon resonance); Lower activation thresholds (functional avidity assays); Enhanced protective capacity (adoptive transfer in mouse models). *Falsification criterion:* If public and private TCRs show equivalent functional parameters, selection is refuted.

#### Prediction 2 (Curation Bias Hypothesis)

If public enrichment is an artifact, prospectively sampled repertoires (unbiased by publication) should yield public fractions matching neutral predictions (∼1.5%). *Falsification criterion:* If unbiased sampling still yields 3% public fraction, curation bias is refuted.

#### Prediction 3 (Convergence Mechanism)

If selection drives convergence, public TCRs should cluster in sequence space around optimal binding motifs. Neutral convergence predicts random (non-clustered) public clone distribution. *Falsification criterion:* Use GLIPH [18] or TCRdist to test for non-random clustering. Random clustering refutes selection.

### Methodological Contributions Beyond Immunology

Our falsificationist framework addresses a general problem in complex systems: distinguishing phenomenological patterns (heavy tails, high diversity) from mechanistic processes (drift vs. selection vs. criticality). Key methodological innovations include:

#### Pre-Validated Null Models

We validate the neutral model against independent benchmarks before comparing to the target dataset. This prevents "strawman" nulls constructed to be easily rejected.

#### Multi-Observable Testing

Single statistics (e.g., power-law exponents) are mechanistically ambiguous. Testing multiple observables with Bonferroni correction increases specificity.

#### Honest Power Reporting

We explicitly calculate and report statistical power, acknowledging when sample sizes are insufficient to detect small effects. This prevents over-interpretation of null results.

#### Effect Size Emphasis

We prioritize Cohen’s d over p-values, quantifying the magnitude of deviations rather than just their statistical significance [25].

These principles generalize to: Antibody repertoire analysis (B-cell convergence); Microbiome ecology (neutral vs. niche-based assembly); Tumor evolution (neutral drift vs. selective sweeps); Neural avalanche dynamics (criticality vs. noise). In each case, rigorous null models—validated independently, tested with multiple observables, and evaluated with honest power—enable mechanistic inference beyond descriptive phenomenology.

### Implications for Vaccine Design and Precision Medicine

Our finding that public clonotypes exceed neutral expectations—while overall diversity remains consistent—has practical implications:

#### Public-First Vaccine Design

If public enrichment reflects convergent selection (Prediction 1 confirmed experimentally), vaccine immunogens should be designed to elicit public TCR responses. This maximizes cross-population coverage and minimizes inter-individual variability.

#### Cautious Biomarker Use

If public enrichment is curation bias (Prediction 2 confirmed), using public TCRs as immune response biomarkers may be misleading. Prospective, unbiased repertoire profiling is essential for clinical validation.

#### Repertoire-Informed Diagnostics

Discriminating neutral from non-neutral repertoire features could improve immune monitoring in cancer immunotherapy, transplant rejection, and autoimmunity. Our framework provides the statistical rigor necessary for such applications.

## Conclusions

We developed and rigorously validated a biologically calibrated neutral null model for TCR repertoire occurrence patterns, enabling falsifiable hypothesis testing in database-aggregated adaptive immunity. Public clonotype frequencies in VDJdb (3.10%) deviate significantly from neutral predictions (1.47%±0.29%, p < 0.001, Cohen’s d = 5.62), inconsistent with neutrality at stringent Bonferroni-corrected thresholds.

Critically, power-law exponents and Shannon entropy remain neutral-consistent, revealing that deviations (if biological) operate specifically on cross-individual convergence rather than overall repertoire diversity. This pattern is consistent with but does not prove epitope-driven functional selection; database curation bias and technical artifacts remain plausible alternatives requiring prospective experimental validation.

Our framework—featuring independent model validation, pre-registered decision criteria, multi-observable testing, and honest effect size reporting—establishes falsificationist methodology for mechanistic inference. The fundamental lesson is epistemological: observing patterns (heavy tails, high diversity) is insufficient for mechanism claims; only explicit, falsifiable null models enable rigorous discrimination between drift, selection, and criticality in complex biological systems.

## Acknowledgments

We thank Universidad Simón Bolívar for institutional support and access to computational resources. We are grateful to the Asociación Colombiana de Inmunología for fostering scientific collaboration. We acknowledge the VDJdb consortium for maintaining a high-quality public database. We thank Dr. Cecilia Fernández Ponce (Universidad de Cádiz, Spain) for valuable discussions on statistical power analysis. We are grateful to our laboratory colleagues at CICV for critical feedback during manuscript preparation.

## Supporting Information

**S1 Fig.** VDJdb Database Composition and Quality Metrics

**S2 Fig.** Pathogen Stratification Analysis: Universality of Public Enrichment

**S3 Fig.** Temporal Stability Analysis (2009-2024): No Systematic Trend

**S4 Fig.** Study Size Independence: No Correlation with Public Fraction

**S5 Fig.** V/J Gene Independence Verification: Naive vs. Experienced Repertoires

**S1 Table.** Sample Size Requirements for Adequate Statistical Power

**S1 Note.** Sensitivity Analysis: Public Clonotype Fraction Threshold Robustness

**S1 Data.** Processed VDJdb Dataset (CSV format)

**S1 Code.** Neutral Model Implementation (Python with documentation)

## References

1. Davis MM, Bjorkman PJ. T-cell antigen receptor genes and T-cell recognition. Nature. 1988;334(6181):395–402. doi:10.1038/334395a0

2. Mason D. A very high level of crossreactivity is an essential feature of the T-cell receptor. Immunol Today. 1998;19(9):395–404. doi:10.1016/s0167-5699(98)01299-7

3. Robins HS, Campregher PV, Srivastava SK, Wacher A, Turtle CJ, Kahsai O, et al. Comprehensive assessment of T-cell receptor beta-chain diversity in alphabeta T cells. Blood. 2009;114(19):4099–4107. doi:10.1182/blood-2009-04-217604

4. Clauset A, Shalizi CR, Newman MEJ. Power-law distributions in empirical data. SIAM Rev. 2009;51(4):661–703. doi:10.1137/070710111

5. Cohen J. Statistical Power Analysis for the Behavioral Sciences. 2nd ed. Hillsdale, NJ: Lawrence Erlbaum Associates; 1988.

6. Hubbell SP. The Unified Neutral Theory of Biodiversity and Biogeography. Princeton, NJ: Princeton University Press; 2001.

7. Bagaev DV, Vroomans RMA, Samir J, Stervbo U, Rius C, Dolton G, et al. VDJdb in 2019: database extension, new analysis infrastructure and a T-cell receptor motif compendium. Nucleic Acids Res. 2020;48(D1):D1057-D1062. doi:10.1093/nar/gkz874

8. Shugay M, Bagaev DV, Zvyagin IV, Vroomans RM, Crawford JC, Dolton G, et al. VDJdb: a curated database of T-cell receptor sequences with known antigen specificity. Nucleic Acids Res. 2018;46(D1):D419–D427. doi:10.1093/nar/gkx760

9. Tickotsky N, Sagiv T, Prilusky J, Shifrut E, Friedman N. McPAS-TCR: a manually curated catalogue of pathology-associated T cell receptor sequences. Bioinformatics. 2017;33(18):2924–2929. doi:10.1093/bioinformatics/btx286

10. Venturi V, Kedzierska K, Turner SJ, Doherty PC, Davenport MP. Methods for comparing the diversity of samples of the T cell receptor repertoire. J Immunol Methods. 2007;321(1-2):182–195. doi:10.1016/j.jim.2007.01.019

11. Venturi V, Quigley MF, Greenaway HY, Ng PC, Ende ZS, McIntosh T, et al. A mechanism for TCR sharing between T cell subsets and individuals revealed by pyrosequencing. J Immunol. 2011;186(7):4285–4294. doi:10.4049/jimmunol.1003898

12. Robins HS, Srivastava SK, Campregher PV, Turtle CJ, Andriesen J, Riddell SR, et al. Overlap and effective size of the human CD8+ T cell receptor repertoire. Sci Transl Med. 2010;2(47):47ra64. doi:10.1126/scitranslmed.3001442

13. Arstila TP, Casrouge A, Baron V, Even J, Kanellopoulos J, Kourilsky P. A direct estimate of the human alphabeta T cell receptor diversity. Science. 1999;286(5441):958–961. doi:10.1126/science.286.5441.958

14. Douek DC, McFarland RD, Keiser PH, Gage EA, Massey JM, Haynes BF, et al. Changes in thymic function with age and during the treatment of HIV infection. Nature. 1998;396(6712):690–695. doi:10.1038/25374

15. Britanova OV, Putintseva EV, Shugay M, Merzlyak EM, Turchaninova MA, Staroverov DB, et al. Age-related decrease in TCR repertoire diversity measured with deep and normalized sequence profiling. J Immunol. 2014;192(6):2689–2698. doi:10.4049/jimmunol.1302064

16. Merkenschlager M, Graf D, Lovatt M, Bommhardt U, Zamoyska R, Fisher AG. How many thymocytes audition for selection? J Exp Med. 1997;186(7):1149–1158. doi:10.1084/jem.186.7.1149

17. Sprent J, Surh CD. Normal T cell homeostasis: the conversion of naive cells into memory-phenotype cells. Nat Immunol. 2011;12(6):478–484. doi:10.1038/ni.2018

18. Glanville J, Huang H, Nau A, Hatton O, Wagar LE, Rubelt F, et al. Identifying specificity groups in the T cell receptor repertoire. Nature. 2017;547(7661):94–98. doi:10.1038/nature22976

19. Dash P, Fiore-Gartland AJ, Hertz T, Wang GC, Sharma S, Souquette A, et al. Quantifiable predictive features define epitope-specific T cell receptor repertoires. Nature. 2017;547(7661):89–93. doi:10.1038/nature22383

20. Emerson RO, DeWitt WS, Vignali M, Gravley J, Hu JK, Osborne EJ, et al. Immunosequencing identifies signatures of cytomegalovirus exposure history and HLA-mediated effects on the T cell repertoire. Nat Genet. 2017;49(5):659–665. doi:10.1038/ng.3822

21. Alstott J, Bullmore E, Plenz D. powerlaw: A Python package for analysis of heavy-tailed distributions. PLoS One. 2014;9(1):e85777. doi:10.1371/journal.pone.0085777

22. Broido AD, Clauset A. Scale-free networks are rare. Nat Commun. 2019;10:1017. doi:10.1038/s41467-019-08746-5

23. Virkar Y, Clauset A. Power-law distributions in binned empirical data. Ann Appl Stat. 2014;8(1):89–119. doi:10.1214/13-AOAS710

24. Button KS, Ioannidis JPA, Mokrysz C, Nosek BA, Flint J, Robinson ESJ, et al. Power failure: why small sample size undermines the reliability of neuroscience. Nat Rev Neurosci. 2013;14(5):365–376. doi:10.1038/nrn3475

25. Lakens D. Calculating and reporting effect sizes to facilitate cumulative science: a practical primer for t-tests and ANOVAs. Front Psychol. 2013;4:863. doi:10.3389/fpsyg.2013.00863

26. Wasserstein RL, Lazar NA. The ASA statement on p-values: context, process, and purpose. Am Stat. 2016;70(2):129–133. doi:10.1080/00031305.2016.1154108

27. Mora T, Walczak AM. How many different clonotypes do immune repertoires contain? Curr Opin Syst Biol. 2016;10:1–7. doi:10.1016/j.coisb.2016.11.001

28. Desponds J, Mora T, Walczak AM. Fluctuating fitness shapes the clone-size distribution of immune repertoires. Proc Natl Acad Sci USA. 2016;113(2):274–279. doi:10.1073/pnas.1512977112

29. Lythe G, Callard RE, Hoare RL, Molina-París C. How many TCR clonotypes does a body maintain? J Theor Biol. 2016;389:214–224. doi:10.1016/j.jtbi.2015.10.016

30. Kimura M. The Neutral Theory of Molecular Evolution. Cambridge: Cambridge University Press; 1983.

31. Elhanati Y, Murugan A, Callan CG Jr, Mora T, Walczak AM. Quantifying selection in immune receptor repertoires. Proc Natl Acad Sci USA. 2014;111(27):9875–9880. doi:10.1073/pnas.1409572111

32. Kared H, Redd AD, Bloch EM, Bonny TS, Sumatoh H, Kairi F, et al. SARS-CoV-2-specific CD8+ T cell responses in convalescent COVID-19 individuals. J Clin Invest. 2021;131(5):e145476. doi:10.1172/JCI145476

33. Nelde A, Bilich T, Heitmann JS, Maringer Y, Salih HR, Roerden M, et al. SARS-CoV-2-derived peptides define heterologous and COVID-19-induced T cell recognition. Nat Immunol. 2021;22(1):74–85. doi:10.1038/s41590-020-00808-x

34. Shomuradova AS, Vagida MS, Sheetikov SA, Zornikova KV, Kiryukhin D, Titov A, et al. SARS-CoV-2 epitopes are recognized by a public and diverse repertoire of human T cell receptors. Immunity. 2020;53(6):1245–1257.e5. doi:10.1016/j.immuni.2020.11.004

35. Ioannidis JPA. Why most published research findings are false. PLoS Med. 2005;2(8):e124. doi:10.1371/journal.pmed.0020124

36. Munafò MR, Nosek BA, Bishop DVM, Button KS, Chambers CD, Percie du Sert N, et al. A manifesto for reproducible science. Nat Hum Behav. 2017;1:0021. doi:10.1038/s41562-016-0021

37. Joglekar AV, Leonard MT, Jeppson JD, Swift M, Li G, Wong S, et al. T cell antigen discovery via signaling and antigen-presenting bifunctional receptors. Nat Methods. 2019;16(2):191–198. doi:10.1038/s41592-018-0304-8

38. Hsieh CL, Goldsmith JA, Schaub JM, DiVenere AM, Kuo HC, Javanmardi K, et al. Structure-based design of prefusion-stabilized SARS-CoV-2 spikes. Science. 2020;369(6510):1501–1505. doi:10.1126/science.abd0826

39. Popper KR. The Logic of Scientific Discovery. London: Routledge; 1959.

40. Platt JR. Strong inference. Science. 1964;146(3642):347-353. doi:10.1126/science.146.3642.347

41. Stumpf MP, Porter MA. Critical truths about power laws. Science. 2012;335(6069):665–666. doi:10.1126/science.1216142

42. Moretti P, Muñoz MA. Griffiths phases and the stretching of criticality in brain networks. Nat Commun. 2013;4:2521. doi:10.1038/ncomms3521

43. Bak P, Tang C, Wiesenfeld K. Self-organized criticality: an explanation of the 1/f noise. Phys Rev Lett. 1987;59(4):381–384. doi:10.1103/PhysRevLett.59.381

44. Zvyagin IV, Minaev I, Pogorelyy MV, Mamedov IZ, Chudakov DM, Shugay M. Distinctive properties of identical twins’ TCR repertoires revealed by high-depth sequencing. Proc Natl Acad Sci USA. 2020;117(6):3009–3018. doi:10.1073/pnas.1913023117

45. Minervina AA, Pogorelyy MV, Komech EA, Kurnikova MA, Shugay M, Chudakov DM. Primary and secondary anti-viral response captured by the dynamics and phenotype of individual T cell clones. Elife. 2020;9:e53704. doi:10.7554/eLife.53704

