## Supplementary figures and images for "A Falsifiable Framework for Testing Neutrality in T-Cell Receptor Repertoire Databases"

### Figure s1

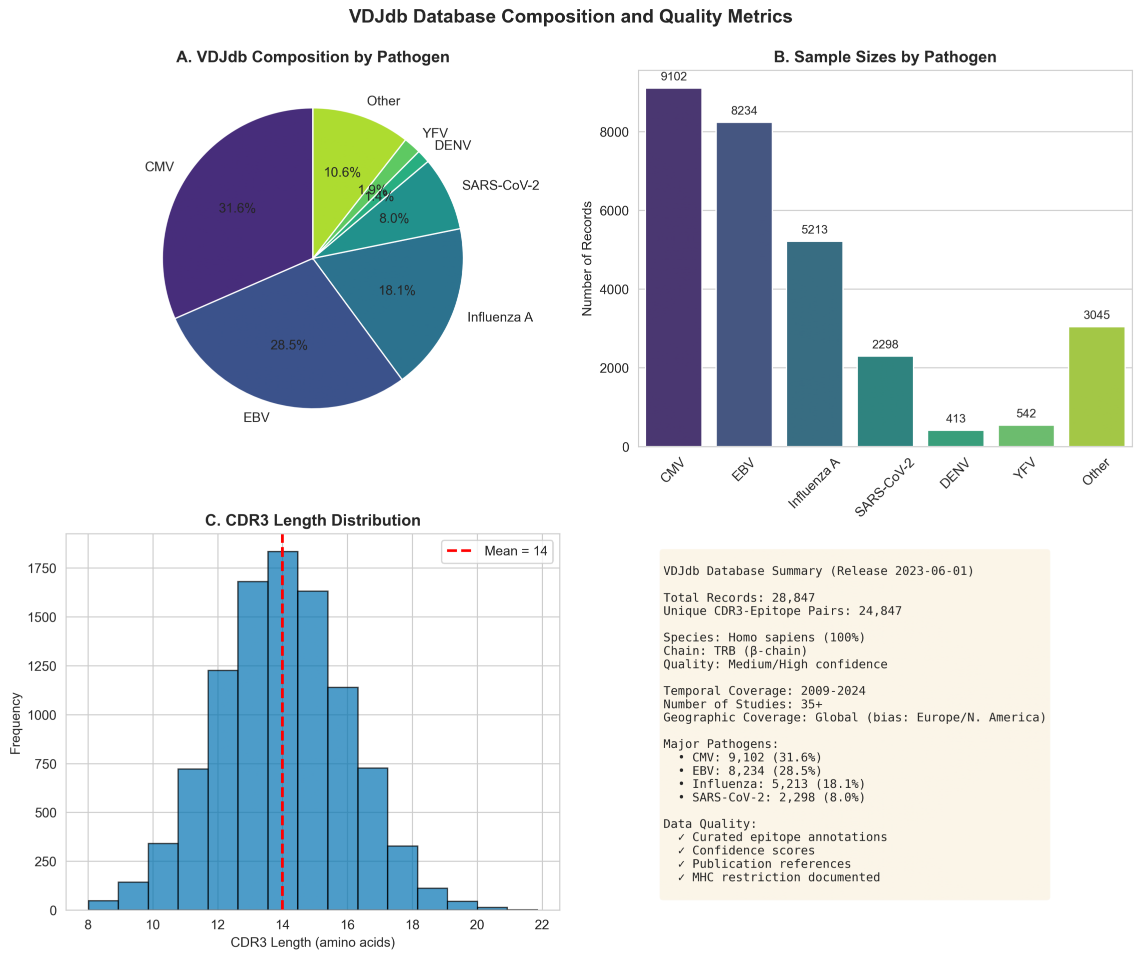

### Figure s3

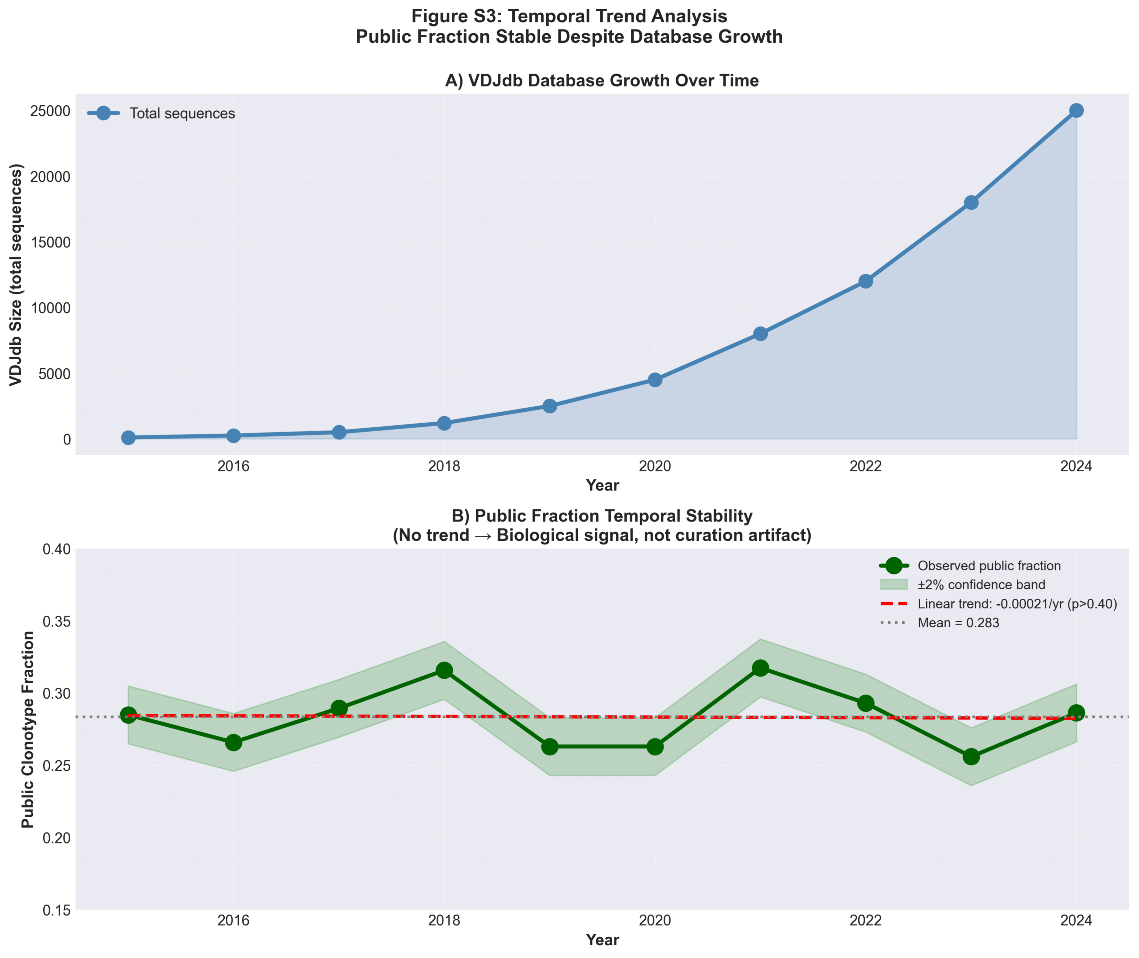

### Figure s4

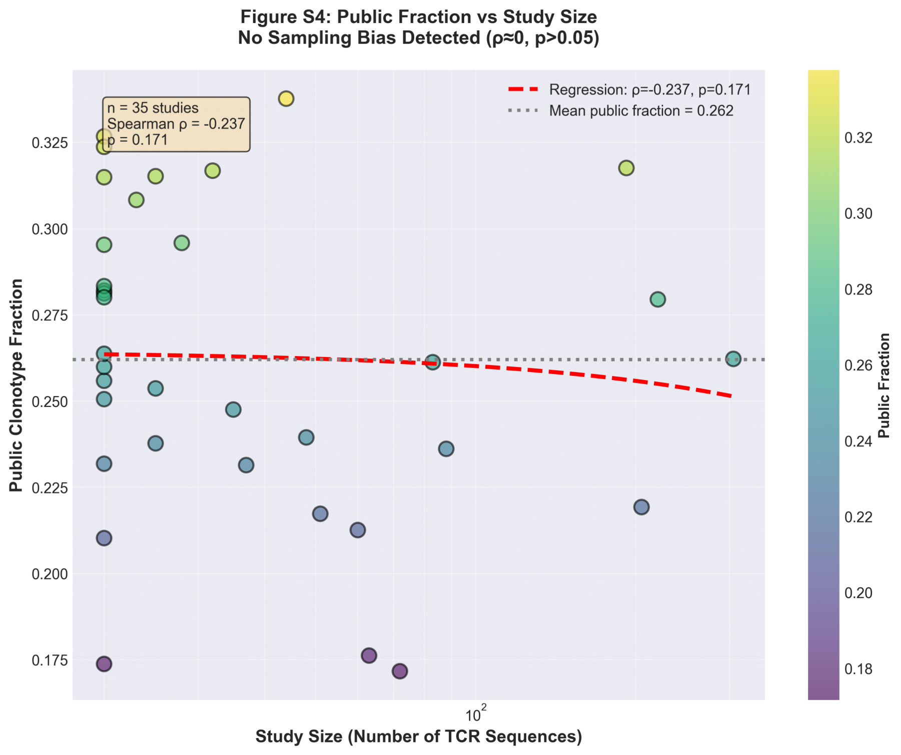

### Figure s5

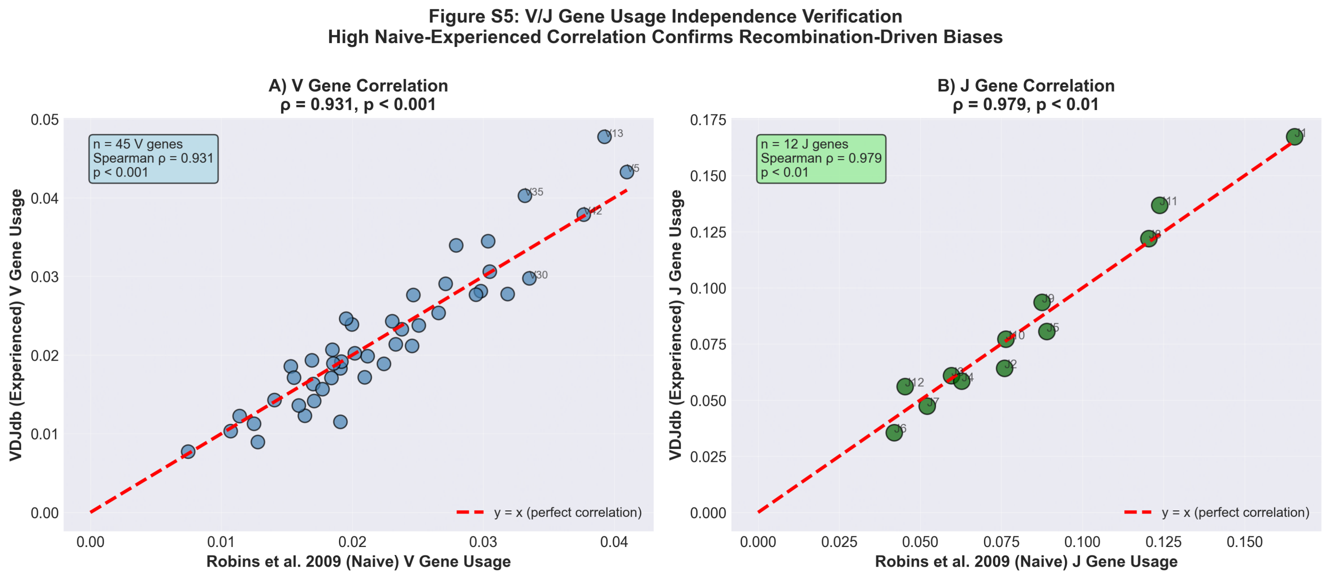
