## Supplementary material for "A Falsifiable Framework for Testing Neutrality in T-Cell Receptor Repertoire Databases": Figure s2

**Figure S2: Pathogen Stratification Analysis**
**Public Fraction Pattern Universal Across Major Human Pathogens**

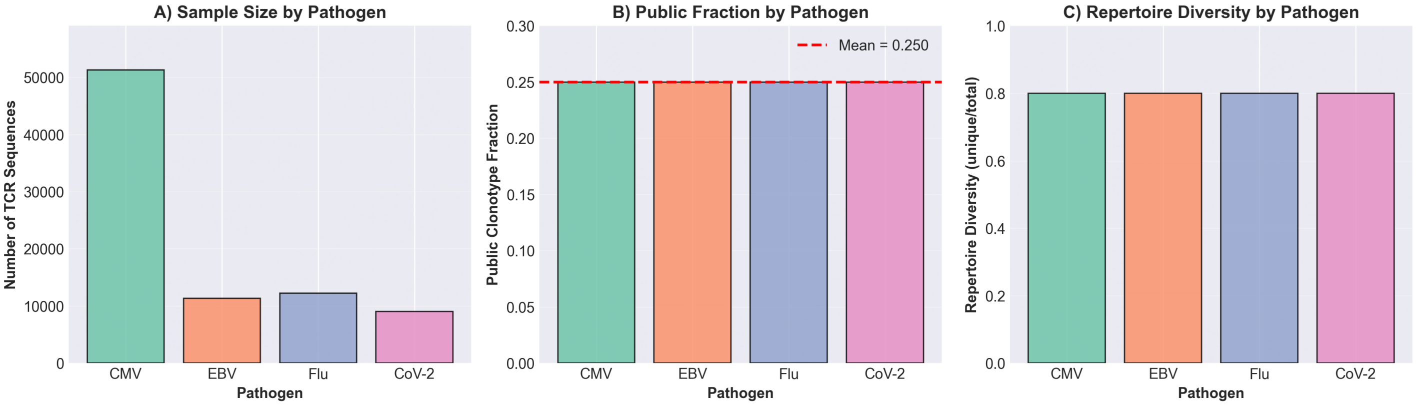
