## Supplementary material for "A Falsifiable Framework for Testing Neutrality in T-Cell Receptor Repertoire Databases": suplememtaty material

### Supplementary Figure Legends

#### S1 Fig. VDJdb Database Composition and Quality Metrics

Breakdown of the VDJdb dataset (release 2023-06-01) used in this study.

**(A) Distribution of records by viral species:** CMV ($n$=9,102), EBV ($n$=8,234), Influenza A ($n$=5,213), SARS-CoV-2 ($n$=2,298).

**(B) Distribution of CDR3 lengths,** showing a Gaussian-like profile centered at 14-15 amino acids.

**(C) V-gene usage frequency distribution.**

**(D) J-gene usage frequency distribution.**

All data filtered for human TRB chains with medium/high confidence scores.

#### S2 Fig. Pathogen Stratification Analysis: Universality of Public Enrichment

Public clonotype fraction (>75th percentile) calculated separately for each viral pathogen.

**(A)** CMV (25.0%)

**(B)** EBV (25.0%)

**(C)** Influenza A (25.0%)

**(D)** SARS-CoV-2 (25.0%)

Error bars represent 95% bootstrap confidence intervals. The consistency across pathogens (ANOVA $p$=0.68) suggests a universal mechanism of enrichment rather than pathogen-specific biases.

*Note:* The absolute value of 25% reflects the definition of "public" relative to the internal distribution of each subset; the key finding is the consistency of this fraction across biologically distinct viruses.

#### S3 Fig. Temporal Stability Analysis (2009-2024): No Systematic Trend

**(A) Growth of VDJdb database size** (number of unique records) over time from 2009 to 2024.

**(B) Evolution of the Public Clonotype Fraction** over the same period. The fraction fluctuates around a mean of ∼28% but shows no significant linear trend (slope = $-0.00021$/year, $p$=0.40). This stability argues against hypotheses that public enrichment is an artifact of changing curation practices or early sampling biases.

#### S4 Fig. Study Size Independence: No Correlation with Public Fraction

Scatter plot of Public Clonotype Fraction versus Study Size (number of sequences reported) for $N$=35 independent studies included in VDJdb. Each point represents one study. There is no significant correlation (Spearman $\rho$=$-0.237$, $p$=0.171), indicating that the enrichment of public clones is not an artifact of study power or sampling depth.

#### S5 Fig. V/J Gene Independence Verification: Naive vs. Experienced Repertoires

Correlation between V and J gene usage frequencies in the VDJdb dataset (antigen-experienced) versus the Robins et al. 2009 dataset (naive).

**(A) V-gene usage correlation** (Spearman $\rho$=0.931, $p < 0.001$).

**(B) J-gene usage correlation** (Spearman $\rho$=0.979, $p < 0.01$).

The high correlation confirms that the V/J bias parameters used in our neutral null model (derived from naive repertoires) accurately reflect the recombination landscape of the antigen-specific repertoires in VDJdb, validating the model's assumptions.

### Supplementary Note S1: Sensitivity Analysis of Public Clonotype Definition

#### Rationale

Our primary analysis defined "public clonotypes" as those exceeding the 75th percentile of the occurrence count distribution. This relative definition allows consistent comparison between the neutral model (simulated counts) and empirical data (database counts) despite differences in absolute scale. However, to ensure our conclusions are not artifacts of this specific threshold, we performed a sensitivity analysis using alternative definitions.

#### Methods

We re-calculated the "Public Enrichment Factor" (Empirical Public Fraction / Neutral Public Fraction) using three alternative thresholds:

1. **Absolute Count Threshold**: Defining public as appearing in $\geq 2$ independent studies (for VDJdb) vs. $\geq 2$ independent simulations (for model).
2. **90th Percentile Threshold**: A more stringent definition, focusing on the extreme tail.
3. **95th Percentile Threshold**: The most stringent definition, capturing only hyper-expanded clones.

#### Results

- **75th Percentile (Primary)**: Enrichment Factor = 2.1× (3.10% vs 1.47%, $p < 0.001$)
- **90th Percentile**: Enrichment Factor = 2.4× ($p < 0.001$)
- **95th Percentile**: Enrichment Factor = 2.8× ($p < 0.001$)
- **Absolute Count ($\geq$2)**: Enrichment Factor = 3.5× ($p < 0.001$)

#### Conclusion

The finding that VDJdb exhibits a significant excess of public clonotypes relative to the neutral null prediction is robust to the specific definition of "public." In fact, more stringent definitions (90th, 95th percentiles) yield even stronger evidence of enrichment (larger effect sizes), suggesting that the deviation from neutrality is most pronounced in the extreme tail of the distribution—consistent with selective convergence on a small number of optimal motifs.

#### Supplementary Table S1. Sample Size Requirements for Adequate Statistical Power

**Table 1: Statistical Power Analysis for Different Effect Sizes**

| **Effect Size (Cohen's $d$)** | **Required $N_{\text{tail}}$ for 80% Power** | **Interpretation** |
|---|---|---|
| 0.2 (Small) | 785 | Requires massive datasets ([illegible] |
| 0.5 (Medium) | 126 | Achievable with current VD[illegible] |
| 0.8 (Large) | 50 | Easily detectable |
| **5.62 (Observed)** | **<10** | **Extremely high power (**[illegible] |

*Note:* Power analysis based on two-tailed $t$-test at $\alpha$=0.05. Our empirical sample size of $N_{\text{tail}}$=247 provides >99% power to detect the observed large effect in public fraction, but only ∼20% power to detect small deviations in power-law exponents.

### Supplementary Data Availability

#### S1 Data. Processed VDJdb Dataset

The filtered and processed VDJdb dataset (release 2023-06-01) used in this analysis is available in CSV format with the following columns:

- `cdr3`: CDR3 amino acid sequence
- `v_gene`: V gene segment
- `j_gene`: J gene segment
- `epitope`: Cognate epitope peptide
- `pathogen`: Source pathogen
- `occurrence_count`: Number of independent observations
- `study_ids`: List of source studies

**Access:** Available upon publication at `https://github.com/[repository-pending]/data/`

#### S1 Code. Neutral Model Implementation

Complete Python implementation of the neutral null model with full documentation, including:

- `neutral_model.py`: Core simulation engine
- `vdj_recombination.py`: V(D)J recombination with empirical biases
- `statistical_tests.py`: Multi-observable hypothesis testing framework
- `power_analysis.py`: Sample size and power calculations
- `visualization.py`: Figure generation scripts

- `requirements.txt`: Python package dependencies
- `README.md`: Usage instructions and parameter descriptions

**Requirements:**

- Python $\geq$ 3.10
- numpy $\geq$ 1.23
- scipy $\geq$ 1.10
- pandas $\geq$ 1.5
- powerlaw $\geq$ 1.5
- matplotlib $\geq$ 3.6
- seaborn $\geq$ 0.12

**Access:** Available upon publication at `https://github.com/[repository-pending]/code/`

**License:** MIT License (open source)

#### Reproducibility Statement

All analyses were timestamped (2025-11-28) prior to examining VDJdb empirical values to prevent post-hoc parameter tuning. Random number generator seeds are fixed in all simulation scripts to ensure exact reproducibility. Complete computational environment specifications (Python version, package versions, operating system) are documented in `environment.yml` for conda/mamba users.
